# Improved Heat Tolerance in Heat-acclimated Mice: The Probable Role of the PD-L1 Pathway

**DOI:** 10.1101/2022.02.20.481185

**Authors:** Minyue Qiu, Yuxin Zhou, Nan Ye, Hongxia Guo, Xiaoyang Zhou, Xiaoyan Ding, Jintao Li

## Abstract

Heat stroke is a life-threatening illness and is related to systematic inflammation-induced multiple organ dysfunction. Available evidence indicates that the severity of the systematic inflammatory response in heat stroke may be related to the changes in immune regulation brought by heat acclimation. However, the mechanisms of heat acclimation are still unclear. Here, we assessed the differences in immunocyte subsets in the spleen and lymph nodes of heat-acclimated and unacclimated mice. A higher frequency of CD4^+^Foxp3^+^ Tregs was observed in heat-acclimated mice. Our results indicated that the improved heat tolerance exhibited during acute heat stress exposure was related to an increased number of Tregs. In heat-acclimated mice, an increase in the number of Tregs was able to mitigate the recruitment of neutrophils, inhibit the activation of neutrophils, and suppress the severity of acute inflammation. Increased differentiation and development of Tregs in peripheral immune organs in heat-acclimated mice might stem from enhanced expression of Foxp3 and PD-L1. Our results strongly suggest that the regulatory function of increased Tregs on neutrophils may be regulated through the PD-1/PD-L1 pathway. The anti-inflammatory effects of Tregs have never been studied in the context of heat stress-induced systemic inflammation. Thus, our results on immunoregulation involving Tregs in heat-acclimated mice might be significant for devising a potential treatment for systemic inflammatory response syndrome and heatstroke.

## INTRODUCTION

Heat stroke is the most devastating form of heat illness and is mainly characterized by a systemic inflammatory response and organ damage^[1, 2]^. Although it has a complicated pathogenesis, an excessive systemic inflammatory response plays an essential role during heat stroke^[3]^. Due to an excessive inflammatory response, an intricate interplay between injury to multiple organs and gut-derived endotoxin translocation, accompanied by dysfunction of the intestinal barrier, always results in high mortality^[4, 5]^.

Heat acclimation is an adaptation to the thermal environment^[6]^. A series of physiological and functional adaptations effectively reduce the negative effects of the thermal environment and the risk of heat stroke^[7, 8]^. Although different methods of heat acclimation have been used to prevent heat stroke for many years, the mechanisms underlying its protective effect in heat stroke-induced injuries remain incompletely understood^[9]^.

The intensity of systemic inflammation determines the severity of injuries to multiple organs to a certain degree during heat stroke^[10, 11]^. In most acute inflammatory responses, the activation of inflammatory cells and release of proinflammatory mediators are regulated by immune cells^[9]^. However, changes in the immune status of heat-acclimated individuals have been insufficiently explored. Whether heat acclimation reduces the severity of systemic inflammation in heat stroke by modulating immune cell functions is unclear.

Regulatory T cells (Tregs), a specialized subset of CD4^+^ T cells, play an essential role in suppressing various types of immune responses, especially in maintaining self-tolerance by suppressing the activation and expansion of autoreactive cells^[12, 13]^. Recent studies suggest that Tregs inhibit the acute inflammatory response^[14, 15]^.

In this study, we established a heat-acclimated (HA) mouse model to test the relationship between the frequencies of Tregs in peripheral lymphoid organs and the protective effect of heat acclimation. In addition, *in vitro* experiments were performed to confirm the underlying mechanism. Our analyses uncover the immunoregulation of improved heat tolerance in HA mice. These insights may provide a new approach for the treatment of heat stroke.

## MATERIALS AND Methods

### Animals

Male C57BL/6 mice (8–12 weeks old; weighing 20–25 g) were obtained from the Laboratory Animal Centre (Army Medical University, Chongqing, China). miR-155^-/-^ mice, Foxp3-GFP mice (C57BL/6 background) and PD-L1^-/-^ mice (BALB/c background) were obtained from the Institute of Immunology (Army Medical University, Chongqing, China). All experimental procedures were approved by the Ethics Committee for Animal Experimentation of the Army Medical University and followed the National Institutes of Health Guide for the Care and Use of Laboratory Animals.

### Heat acclimation model and heat stress exposure protocol

Mice were adapted to standard plastic cages for 5 days and then divided into two groups: the unacclimated group, maintained at an ambient temperature (22 ± 2 °C), and the HA group, maintained in a climatic chamber at 35 ± 1 °C for 4 weeks (21). The survival rate and survival time of mice exposed to 150 min of heat stress (HS) in a 43 ± 1 °C climatic chamber were recorded. Tissues were harvested from surviving mice after 150 min of acute HS.

### Flow cytometry analysis

Lymph nodes and spleens from mice were prepared into single cell suspensions for flow cytometry analysis. Briefly, lymph nodes and spleens were disaggregated through 100-μm sieves (BD, USA). Red blood cells were lysed (spleens only), centrifuged (300 g, 5 min, room temperature) and resuspended in 4°C PBS containing 2% FBS. Cells were counted and adjusted to 10^6^ cells/ml and then incubated with the appropriate monoclonal antibody for 30 min. All antibodys were purchased from Ebioscience (USA): anti-CD3(FITC), anti-CD11b(PE), anti-CD11c(APC), anti-NK1.1(Percp-cy5.5), anti-CD25(APC), anti-CD3(FITC), anti-CD4(PE), anti-CD3(PE-Cy5), anti-CD3(PE-Cy7), anti-CD8(FITC), anti-CD4(FITC). The frequency of neutrophils in the peripheral blood of HA mice and unacclimated mice was analyzed individually at 15 min, 30 min, 45 min and 60 min after being put into a 43°C climatic chamber. Venous blood (20 µl) from the tails was collected and mixed into 1 ml PBS. Then, the cells were centrifuged (11000 g, 15 s, room temperature). Red blood cells were lysed, and residual cells were incubated with the appropriate monoclonal antibody for 30 min. Anti-CD11b (PE) and anti-Gr1 (APC) were purchased from eBioscience (USA). The frequencies of immunocyte subsets and neutrophils were detected by a FACSCalibur (BD, USA).

### Histology

Lungs and livers from mice were fixed in 4% paraformaldehyde for 24 hours. The tissues were trimmed and dehydrated in an automation-tissue-dehydrating machine, and then the prepared samples were embedded in paraffin and sliced into 6-mm-thick sections. After staining with hematoxylin and eosin (HE) by conventional methods, the sections were observed with an Olympus microscope. Damage to the lungs and livers was scored by a colleague who was specialized in this type of job and blinded to the treatments. Scoring of liver sections was based on Suzuki scoring, involving (a) the presence and severity of sinusoidal congestion (0: none, 1: minimal, 2: mild, 3: moderate, 4: severe), (b) necrosis of parenchymal cells (0: none, 1: single cell necrosis, 2:< 30%, 3:30%–60%, 4:> 60%) and (c) cytoplasmic vacuolization (0: none, 1: minimal, 2: mild, 3: moderate, 4: severe) ^[16].^ Scoring of lung sections included (a) alveolar edema, interstitial edema, (b) alveolar structural disturbance and (c) inflammatory infiltration. Each criterion of lung sections was scored 0: none, 1: mild; 2: moderate and 3: severe ^[17]^. The score of each tissue was calculated as the sum scores of each criterion, and at least five microscopic areas were assessed in each section^[16, 17]^.

### Quantitative real-time PCR

Total RNA was extracted from spleen and liver tissues of HA and unacclimated mice using TRIzol reagent (Invitrogen, USA), and the concentration of total RNA was measured by an Epoch Multi-volume Spectrophotometer (BioTek, USA). These RNA samples were reverse transcribed into cDNA using a PrimeScript RT reagent Kit (TaKaRa, Japan), and quantitative real-time PCR (q-PCR) was carried out using SYBR Premix Ex Taq (TaKaRa, Japan). Q-PCR was carried out in a Bio-Rad CFX96 system. The data were analyzed using the 2^−ΔΔCT^ method^[18]^. Each experiment was conducted at least in triplicate.

### Immunohistochemistry

Immunohistochemistry was performed on paraffin-embedded sections of spleen tissues. Slides were deparaffinized and rehydrated in xylene and graded ethanol. Peroxidase was inactivated with 3% hydrogen peroxide for 10 min. Then, the slides were incubated with primary antibodies against PD-L1 (Cell Signaling Technology) and Foxp-3 (Cell Signaling Technology) at 4 °C overnight. The sections were then incubated with a secondary anti-rabbit HRP antibody (Zhongshan Golden Bridge Biotechnology, Beijing, China) for 30 min. Visualization was carried out by using DAB chromogen buffer. Before being sealed by cover slips and residence, all sections were counterstained with hematoxylin for 30 s and dried in a drying oven.

### Sorting of Tregs

Tregs from the lymph nodes and spleen of Foxp3-GFP mice or PD-L1^-/-^ mice were isolated using a FACS Aria II system (BD, USA). Briefly, the lymph nodes and spleen were disaggregated through a 100 μm sieve (BD, USA) with phosphate-buffered saline (PBS). Next, the cell suspension was centrifuged (300 × *g*, 5 min, 25 °C), and the cells were resuspended in cold (4 °C) PBS containing 2% fetal bovine serum (FBS). Tregs of Foxp3-GFP mice were sorted directly based on green fluorescent protein fluorescence, whereas Tregs of PD-L1^-/-^ mice were stained with antibodies (anti-CD4:PE and anti-CD25:APC) before sorting. The number of sorted cells was adjusted to 10^7^ cells/ml in PBS (4 °C) containing 2% FBS.

### Adoptive transfer of Tregs

Sorted Treg suspension (Treg transfer group) or PBS (PBS group) was injected into unacclimated mice via the tail vein (100 µl/mouse). An acute HS exposure test was performed 24 h later. The survival rate and survival time of mice were recorded during 150 min of acute HS exposure. Organs were obtained from surviving mice after 150 min of exposure to acute HS.

### Injection of recombinant PD-L1 protein

Recombinant mouse PD-L1 protein dissolved in PBS (50 μg/ml) was injected into unacclimated mice via the tail vein (100 μl/mouse). An acute HS exposure test was performed 3 h later. The survival rate and survival time of mice were recorded during 150 min of acute HS exposure.

### Coculture of Tregs and neutrophils

Tregs were sorted as mentioned above. Then, Tregs were incubated in mixed 1640 medium containing 10% FBS, 50 ng/ml phorbol-12-myristate-13-acetate (PMA) and 200 IU/ml interleukin-2 (IL-2) in a 37°C incubator. After 72 h, the Trigs were collected and resuspended in pure 1640 medium. Neutrophils were isolated from the bone marrow of 8- to 12-week-old C57/BL6 mice via the protocol of Swamydas *et al*.^[19]^. Then, Tregs (6Swa^4^ per well) were incubated with transforming growth factor-medium antibodies, interleukin-10 (IL-10) antibodies, programmed death 1-ligand 1 (PD-L1) antibodies or cytotoxic T-lymphocyte antigen 4 (CTLA-4) antibodies in 24-well culture plates for 1 hour before neutrophil (6ed ^4^ per well) addition. After coculture in a 37°C incubator for 30 min, a 24-well culture plate containing Tregs and neutrophils was moved into a 41°C incubator. After coculture for another 20 min in a 41°C incubator, culture supernatants were collected. All monoclonal antibodies and recombinant mouse CD274 protein were purchased from R&D Systems (USA).

### Myeloperoxidase ELISA

Myeloperoxidase (MPO) concentrations in Tregs and neutrophil culture supernatants were analyzed using an MPO (mouse) ELISA kit (BOSTER Biological Technology, Wuhan, China) according to the manufacturer’s instructions.

### Statistical analyses

All statistical analyses were performed using GraphPad Prism 5 (GraphPad Software Inc., USA). Data are presented as the mean ± SEM. *P* values were calculated by conducting one-sample *t*-test, bivariate, and survival analysis (Kaplan–Meier method) or one-way ANOVA.

## RESULTS

### Frequency of CD4^+^Foxp3^+^ Treg cells is increased in HA mice

Before evaluating the changes in immunocyte subsets in the peripheral immune organs of HA mice, the protective effect of heat acclimation was assessed via the HS exposure experiment. The mortality rates of HA mice and unacclimated mice during the HS experiment were 33.3% and 61.1%, respectively (*p* < 0.05; Supplementary Figure 1A). Organs, such as the liver, lung, gut, and kidney, retrieved from surviving mice after 150 min of HS in the unacclimated group showed different degrees of increased swelling and blood clotting compared with those from the HA group (Supplementary Figure 1B). Cell suspensions prepared from the lymph nodes and spleen of mice were subjected to fluorochrome staining with monoclonal antibodies. No significant differences in the CD3^+^/CD19^+^ cell ratios were noted between HA and unacclimated mice (*p* > 0.05; Supplementary Figure 2A-D). In the spleen and lymph node suspensions of HA mice, the percentages of CD4^+^ T cells and CD8^+^ T cells were similar to those in the suspensions from unacclimated mice (*p* > 0.05; Supplementary Figure 2E, F). Remarkably, the percentage of CD4^+^Foxp3^+^ cells in the cell suspensions of HA mice was significantly increased compared with that in unacclimated mice (*p* < 0.05; Figure 1A–D). No significant differences in the frequencies of CD11b^+^ macrophages, CD11c + dendritic cells, or NK1.1^+^ cells were observed between the HA and unacclimated mice (*p* > 0.05; Supplementary Figure 3A–D).

**Figure 1.**
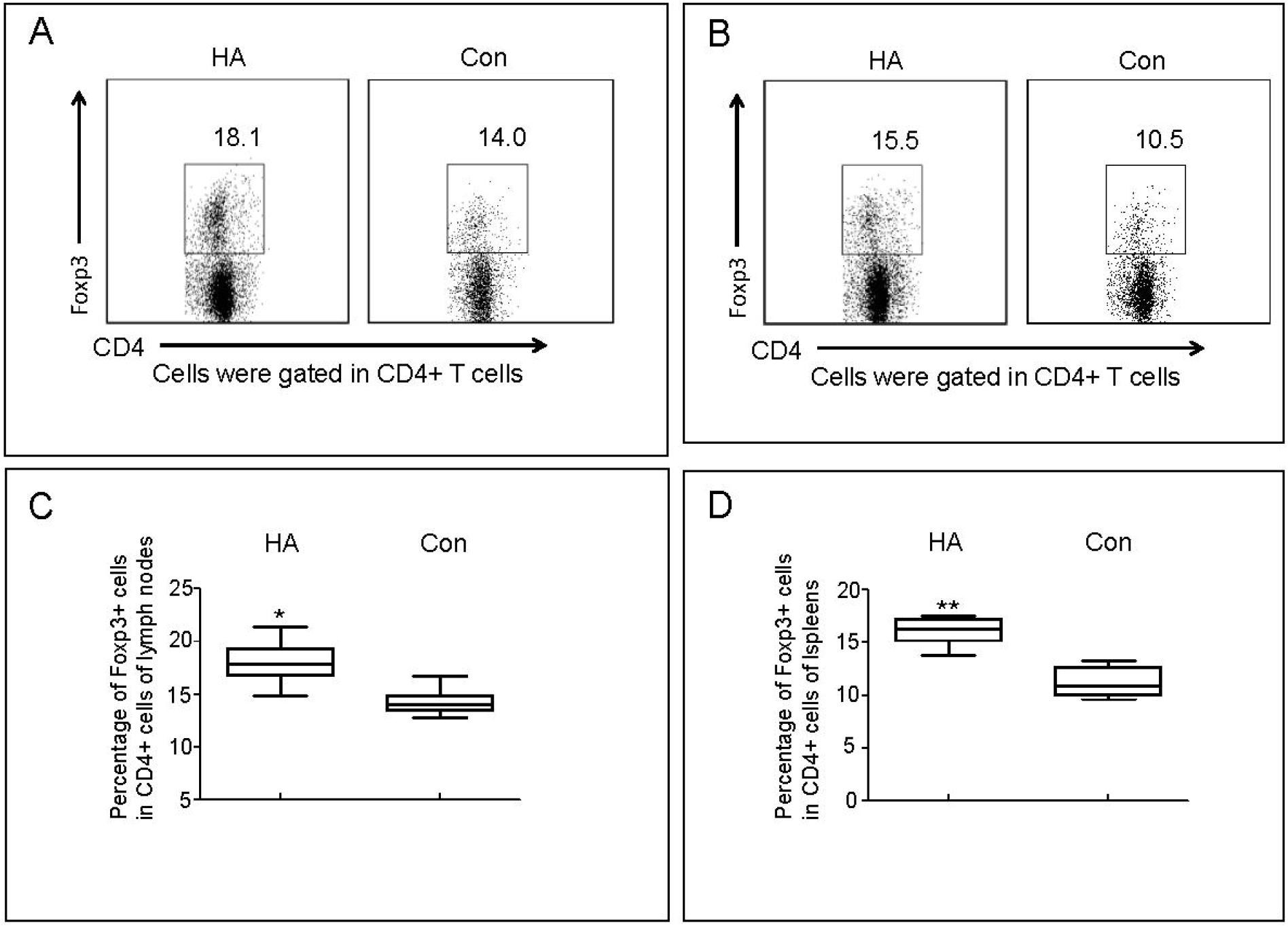
Changes in the frequency of regulatory T cells (Tregs) in the lymph nodes and spleen of BALB/c mice after 4 weeks of heat acclimation. Tissues were harvested from mice in the heat-acclimated (HA) and nonacclimated control (Con) groups. CD4^+^Foxp3^+^ cells were gated from CD4^+^ cells of the lymph nodes (**A**) and spleen (**B**), and then the difference in the frequencies of CD4^+^Foxp3^+^ cells from lymph nodes (**C**) and spleen (**D**) were assessed. Number of mice: unacclimated control, n = 10; heat acclimated, n = 10. \**p* < 0.05, \*\**p* < 0.01 versus the Con group.

### Increased Tregs in HA mice regulate the elevation of neutrophils during acute heat stress

We assessed the frequencies of neutrophils in the peripheral blood of HA and nonacclimated mice during acute HS exposure. The trends at 15, 30, 45, and 60 min are shown in Supplementary Figure 4A. The results suggested a slow elevation in the neutrophil percentage in the peripheral blood of HA mice compared with that in unacclimated mice (*p* < 0.01, Supplementary Figure 4B). The percentage of CD4^+^Foxp3^+^ Tregs in the spleen of HA and nonacclimated mice was reduced after 150 min of acute HS (*p* < 0.05, Figure 2A, C). The induction of higher Tregs might play a pivotal role in protection against HS. Furthermore, the neutrophil frequency in the spleen was significantly decreased in HA mice compared with that in unacclimated mice after HS (*p* < 0.05, Figure 2B, D), and the neutrophil frequency before HS was not significantly different between the HA and unacclimated groups (data not shown).

**Figure 2.**
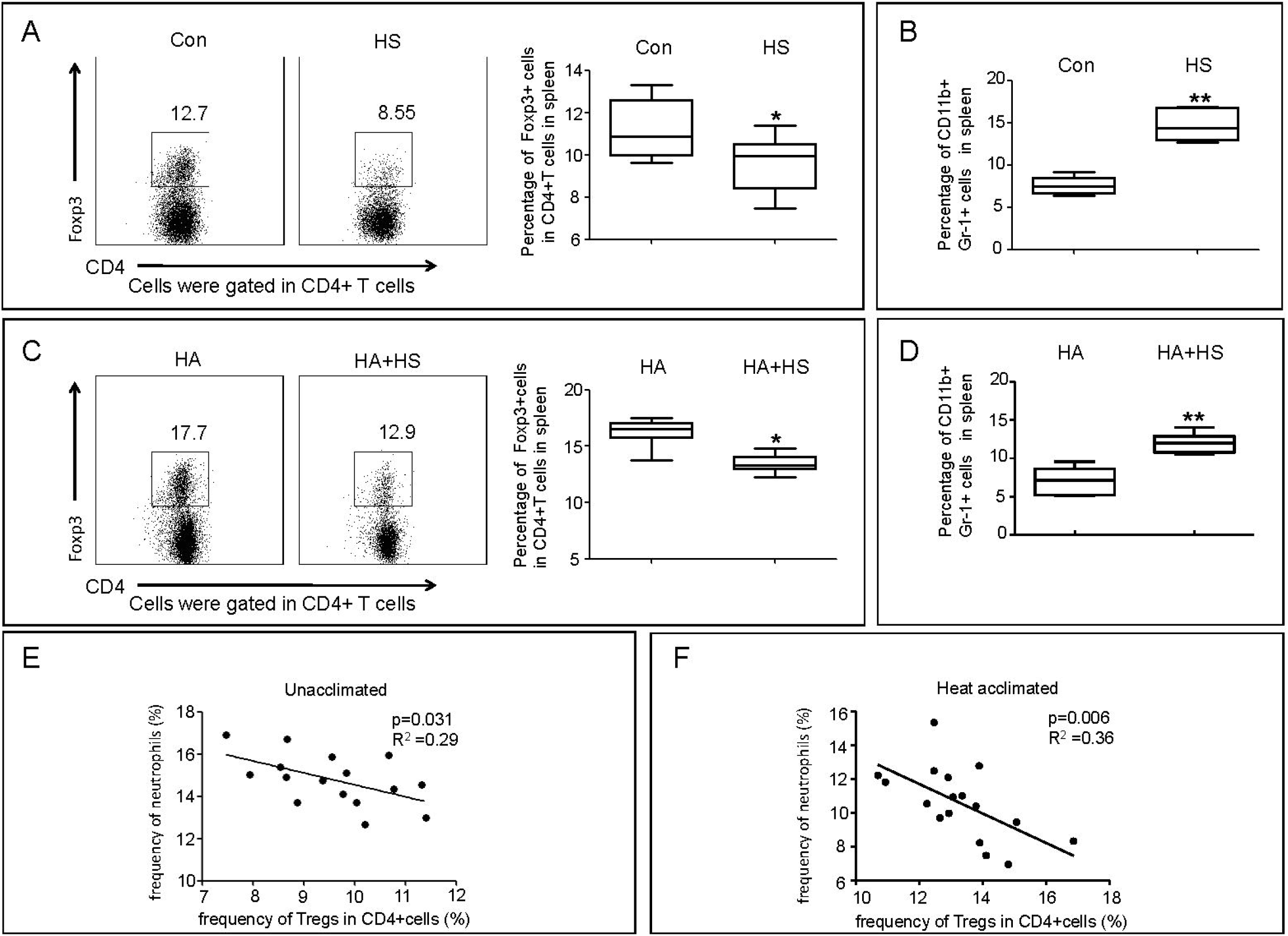
Correlation of regulatory T cells (Tregs) and neutrophils in the spleen of mice in the heat acclimated (HA) and unacclimated control (Con) groups. Unacclimated control (Con) mice and unacclimated mice exposed to 150 min of heat stress (HS) were sacrificed for harvesting spleen cells, and the frequencies of CD4^+^Foxp3^+^ (**A**) and Gr-1^+^CD11b^+^ (**C**) cells were compared. HA mice and HA mice exposed to 150 min of HS (HA+ HS) were sacrificed to harvest spleen cells, and then the frequencies of CD4^+^Foxp3^+^ (**B**) and Gr-1+CD11b+ (**D**) cells were compared. In addition, a correlation analysis was carried out between CD4^+^Foxp3^+^ and Gr-1^+^CD11b^+^ cells in the spleen of mice in the HS (**E**) and HA + HS (**F**) groups. Number of mice: Con group, n = 10; HA group, n = 10; HS group, n = 16; HA + HS group, n = 17. \**p* < 0.05, \*\**p* < 0.01 versus the respective Con group. ^*^*p* < 0.05, ^**^*p* < 0.01 versus the HA group.

These results might be due to the increased suppressive function of Tregs. To determine whether the mitigated recruitment of neutrophils in HA mice was related to increased Tregs, the frequencies of neutrophils and Tregs in the spleen of HA and unacclimated mice were assessed after 150 min of acute HS, after which a bivariate correlation analysis was performed. As shown in Figure 2E and F, in control group mice, a higher frequency of Tregs was accompanied by a lower frequency of neutrophils in the spleen. The highly negative correlation observed indicates that milder neutrophil infiltration in HA mice may be due to higher Treg levels.

### Heat tolerance of mice can be affected by changes in Treg frequencies

To determine whether the improved heat tolerance in HA mice is related to increased Tregs, an adoptive transfer experiment was performed. The purity of sorted Tregs from the spleen and lymph nodes of acclimated mice was assessed (Supplementary Figure 5A), after which they were injected into the unacclimated mice. The effects of transferred Tregs were assessed after HS. The survival of mice in the Treg-transferred group was 83.3% compared with a significantly lower survival (41.7%) in the PBS-transferred group (*p* < 0.05; Figure 3A). The lungs and livers obtained from surviving mice were fixed, embedded, and sliced for hematoxylin and eosin (HE) staining. No obvious abnormalities were observed in the control group, whereas pathological changes were visible in the PBS + HS and Treg transfer + HS groups. However, focal necrosis and infiltration of inflammatory cells in the livers of mice in the PBS + HS group were more severe than those in the Treg transfer + HS group. More obvious vascular engorgement and pulmonary septum broadening were also observed in the PBS + HS group (Supplementary Figure 5B). Compared with the control mouse group, the PBS + HS group also had a significantly higher pathological score, which was attenuated in the Treg transfer + HS group (*p* < 0.05, Figure 3B–C).

**Figure 3.**
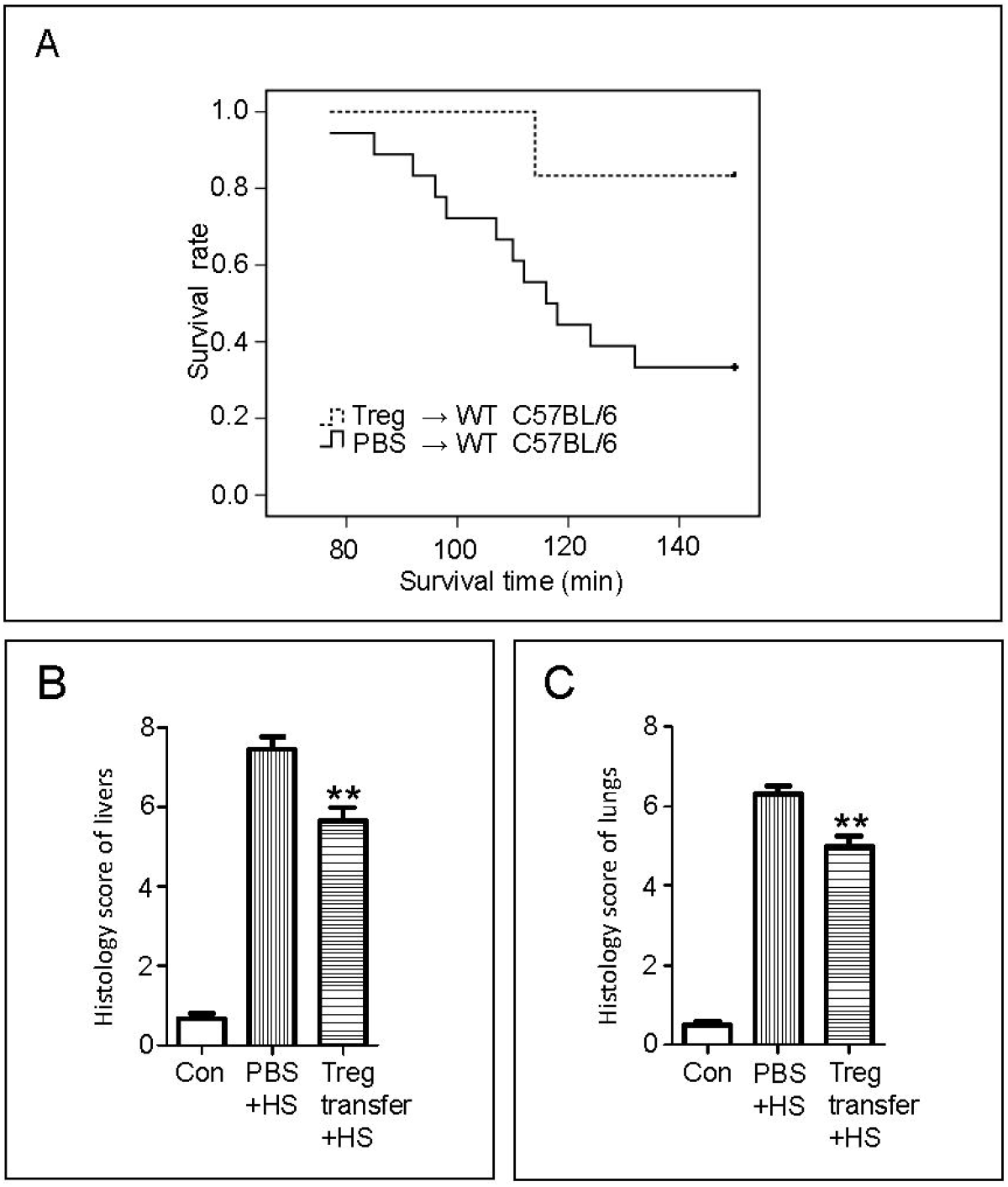
Effect of transferring regulatory T cells (Tregs) on HS exposure. **A**, Treg suspension or phosphate-buffered saline (PBS) was injected into the circulation of unacclimated mice, and the survival rate of mice in the Treg-transferred and PBS groups during exposure for 150 min to HS was recorded. Number of mice: Treg-transferred group, n = 6; PBS group, n = 18. **B**, Liver tissue from mice in the Treg-transferred and PBS groups was harvested after 150 min of HS exposure. In addition, the livers from mice not subjected to any treatment were harvested and used as a control group (Con). Damage to livers was scored by a specialist who was blinded to the treatments. Scoring of liver sections was based on the Suzuki scoring method, involving (a) the presence and severity of sinusoidal congestion (0: none; 1: minimal; 2: mild; 3: moderate; 4: severe), (b) necrosis of parenchymal cells (0: none; 1: single cell necrosis; 2: <30%; 3: 30%–60%; 4: >60%), and (c) cytoplasmic vacuolization (0: none; 1: minimal; 2: mild; 3: moderate; 4: severe). Number of mice: Treg-transferred group, n = 5; PBS group, n = 6; Con group, n = 4. **C**, Lungs from mice in the Treg-transferred and PBS groups were harvested after 150 min of HS exposure. Lungs from mice not subjected to any treatment were harvested and used as a control group (Con). Lung tissue sections stained with hematoxylin and eosin were scored using the lung injury score. Scoring of lung sections included (a) alveolar and/or interstitial edema, (b) alveolar structural disturbance, and (c) inflammatory infiltration. Each criterion for lung sections was scored on a scale of 0–3 (0: none; 1: mild; 2: moderate; 3: severe). Number of mice: Treg-transferred group, n = 5; PBS group, n = 6; Con group, n = 4. \*\**p* < 0.01 versus the respective PBS + HS group.

We tested the mortality of miR-155^-/-^ mice exposed for 150 min to acute HS. miR-155^-/-^ mice tended to die sooner than wild-type (WT; C57BL/6) mice; all 12 miR-155^-/-^ mice died during the 150 min experiment (*p* < 0.05, Figure 4A). The percentage of CD4^+^CD25^+^ Tregs in CD4^+^ T cells was confirmed (*p* < 0.05, Supplementary Figure 6A). Both miR-155^-/-^ and WT mice were placed in a 43 ± 1 °C climatic chamber for 80 min to determine the histology and pathological scores (Supplementary Figure 6B). Liver and lung tissues in miR-155^-/-^ mice exhibited more severe damage, which was associated with a higher pathological score (*p* < 0.01, Figure 4B, C).

**Figure 4.**
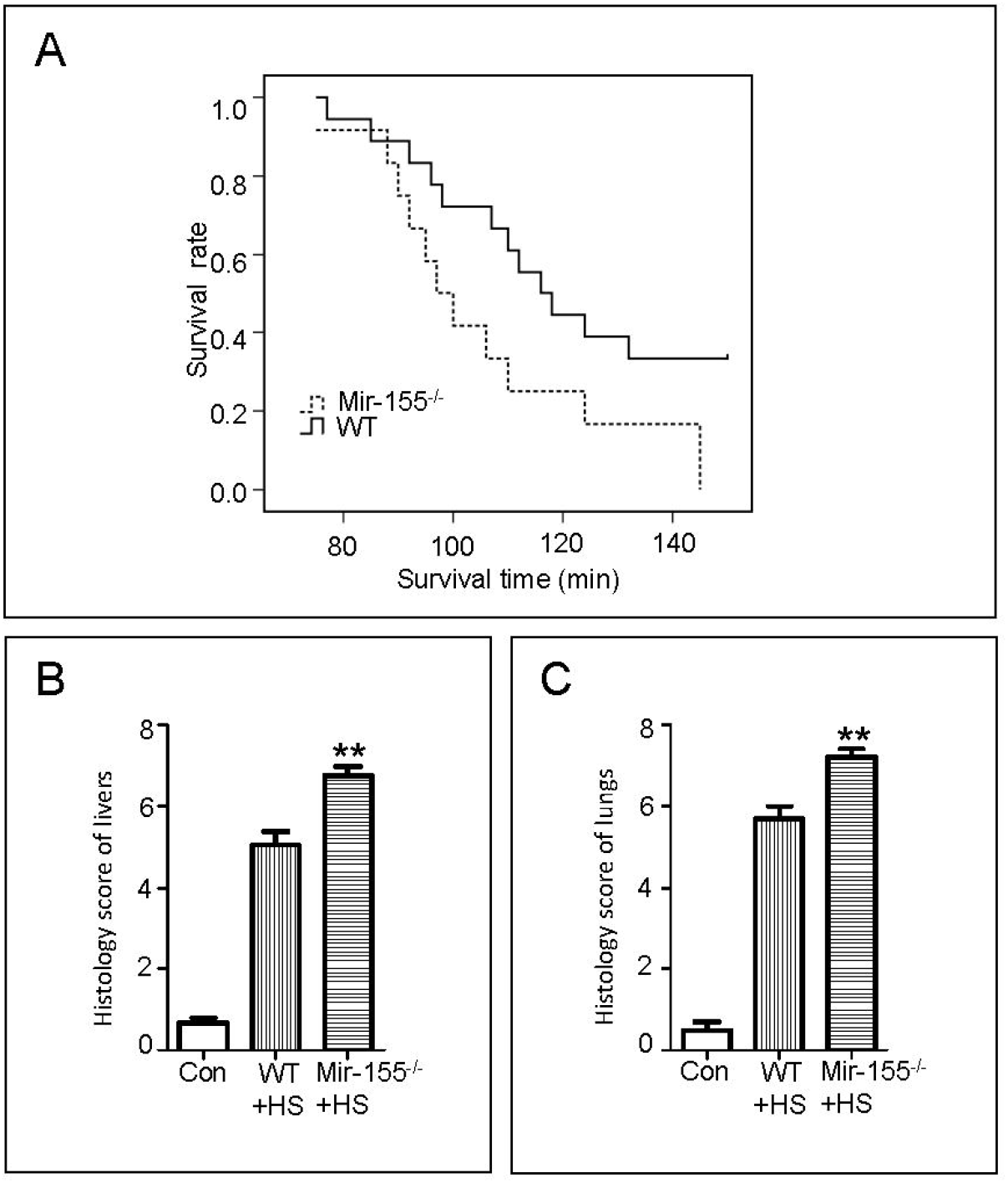
Survival and tissue damage in microRNA (miR)-155^-/-^ mice during heat stress (HS) exposure. **A**, The survival rate of miR-155^-/-^ and wild-type (WT) mice during 150 min of HS exposure was recorded. Number of mice: miR-155^-/-^ mice, n = 12; WT, n = 18. **B**, Lung tissue from miR-155^-/-^ (miR-155^-/-^ + HS) and WT (WT + HS) mice was harvested after 80 min of HS exposure. Lungs from mice not subjected to any treatment were harvested and used as a control group (Con). Lung tissue sections stained with hematoxylin and eosin were scored using the lung injury score. Number of mice: miR-155^-/-^ + HS group, n = 6; WT + HS group, n = 6, Con group, n = 4. **C**, Livers from mice in the Treg-transferred and PBS groups were harvested after 150 min of HS exposure. The livers from mice not subjected to any treatment were harvested and used as a control group (Con). Liver tissue sections stained with hematoxylin and eosin were scored using the liver injury score. Number of mice: miR-155^-/-^ + HS group, n = 6; WT + HS group, n = 6; Con group, n = 4. \*\**p* < 0.01 versus the respective WT + HS group.

### Tregs inhibit the activation of neutrophils through the PD-1/PD-L1 pathway

To explore the mechanism of the anti-inflammatory effect of increased Tregs in HA mice under heat stress, we assessed the changes in the expression of CTLA-4, GATA3, PD-1, PD-L1, and Foxp3 mRNAs in the liver and spleen. No significant differences were found in the expression of CTLA-4 and GATA3 mRNAs. However, the expression of PD-1, PD-L1, and Foxp3 mRNAs was significantly increased in HA mice (Supplementary Figure 7A, B). Immunohistochemistry of the spleen sections showed similar results. The number of Foxp3-positive cells exhibited a slight decrease after HS exposure in both HA and nonacclimated mice. However, the number of these cells increased significantly in the spleen of the HA group compared with that in the control group (Supplementary Figure 8A, B). The number of PD-L1-positive cells in the spleen of the HA group was also greater than that in the control group (Supplementary Figure 8C–D).

To further clarify the suppression of neutrophil recruitment during acute HS, Tregs and neutrophils were cocultured. The purity of the enriched neutrophils and Tregs is shown in Supplementary Figure 9A. Lower MPO levels were observed in the neutrophil and Treg coculture group than in the neutrophil group, and this decrease was abrogated upon pretreatment of Tregs with PD-L1 antibody but not with CTLA-4, PD-L1, TGF-β, or IL-10 antibody (Figure 5A). In view of this finding, we examined the effect of preincubation of neutrophils with different concentrations of recombinant PD-L1 protein; neutrophil-derived MPO levels were reduced in a dose-dependent manner, and the inhibitory effect of Tregs was reversed when neutrophils were preincubated with the PD-1 antibody (Supplementary Figure 9B, C). The mortality upon HS exposure was also significantly lower in the PD-L1 protein-injected group than in the PBS-injected group (6 of 12 mice died in the PBS-injected group, and 1 of 10 mice died in the PD-L1 protein-injected group; *p* < 0.05) (Figure 5B). Remarkably, the survival rate of unacclimated WT (BALB/c background) mice did not significantly increase after injection with Tregs from PD-L1^-/-^ mice.

**Figure 5.**
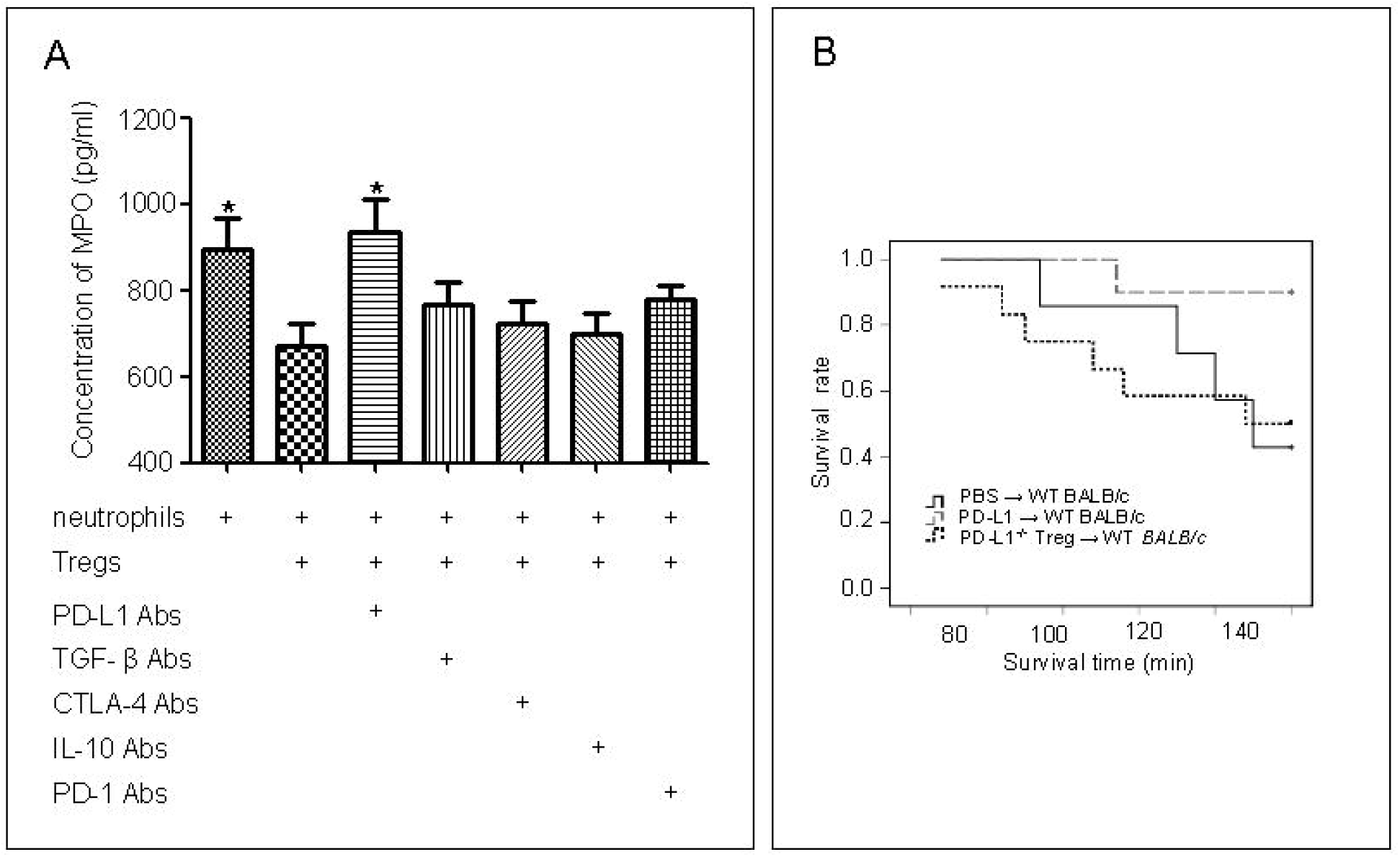
Effect of PD-L1 on heat stress (HS) exposure *in vivo* and *in vitro*. **A**, Regulatory T cells (Tregs) were sorted and incubated in RPMI 1640 medium containing 10% fetal bovine serum, 50 ng/mL phorbol-12-myristate-13-acetate, and 200 IU/mL interleukin-2 in a 37 °C incubator. After 72 h, Tregs were collected and resuspended in RPMI 1640 medium. Neutrophils were isolated from the bone marrow of 8- to 12-week-old C57BL/6 mice via the protocol described by Swamydas et al. (24). Thereafter, Tregs (6 × 10^4^ per well) were incubated with antibodies against transforming growth factor-β (TGF-β), interleukin-10 (IL-10), programmed death 1-ligand 1 (PD-L1), or cytotoxic T-lymphocyte antigen 4 (CTLA-4) in a 24-well culture plate for 1 h before adding neutrophils (6 × 10^4^ per well). After coculture in a 37 °C incubator for 30 min, the 24-well culture plate containing Tregs and neutrophils was moved to a 41 °C incubator. After coculture for an additional 20 min in a 41 °C incubator, the culture supernatants were collected. The concentration of myeloperoxidase (MPO) in the supernatant in the different groups was evaluated. Number of repeated wells: n = 6. \**p* < 0.05 versus the neutrophils + Tregs group. **B**, Recombinant mouse PD-L1 protein, Tregs from PD-L1^-/-^ mice, or PBS was injected into the circulation of wild-type (WT) mice, and the survival rate during 150 min of HS was recorded. Number of mice: PD-L1 protein-injected mice, n = 10; transfer of Tregs from PD-L1^-/-^ mice, n = 7; PBS-injected mice, n = 12.

## DISCUSSION

The main physiological changes associated with HA involve lower core temperature and heart rate and a higher rate of perspiration with reduced minerals in sweat in HS and exercise^[20-22]^. The detailed mechanisms for these changes remain unclear. Studies at the cellular level have mostly focused on the cytoprotection rendered by increased HSF-1, HSP72, and HSP90 levels^[1, 20, 23-25]^. Very little is known about the changes in immunoregulation caused by heat acclimation.

Our primary goal was to assess the changes in the immunocytes in HA mice and to determine whether altered immune regulatory responses contribute to the protective effect in HS. In our experiments, increased frequencies of CD4^+^Foxp3^+^ Tregs were detected in the spleen and lymph nodes after 4 weeks of HA. In a previous study, HA did not result in a significant change in the percentage of CD4^+^ and CD8^+^ T cells in the blood^[26]^. We also did not observe any significant change in the percentage of CD4^+^ and CD8^+^ T cells in the spleen and lymph nodes of HA mice. We found no differences in the frequencies of CD11b^+^, CD11c^+^, CD3^+^, and NK1.1^+^ cells between the HA and unacclimated mice.

Tregs play a pivotal role in modulating the immune system, suppressing inflammatory responses, and limiting tissue damage^[27]^. Lewkowicz et al. indicated that most diseases associated with defective Treg function can be linked to exacerbated neutrophil activity^[28-30]^. As a critical component of innate immunity, neutrophils respond to both bacterial infections and sterile injuries^[30, 31]^. Although they play a protective role in host defense against microorganisms, excessive neutrophil infiltration can cause an overt inflammatory response and severe tissue damage. In a previous study, Treg-induced inhibition of neutrophils was demonstrated in a lipopolysaccharide-stimulated animal model^[32]^. However, the anti-inflammatory effects of Tregs have never been studied in the context of HS-induced systemic inflammation. Based on previous findings and our observation of stably increased Tregs in the spleen and lymph nodes of HA mice, we speculate that heat tolerance induced by HA is related to the inhibition of neutrophils by Tregs. A more rapid elevation in neutrophils was observed in the peripheral blood of unacclimated mice at different phases. The data show that neutrophils may be inhibited by increased Tregs during HS exposure, implying that the development of systemic inflammatory response syndrome could be slower in HA mice than in unacclimated mice under the same HS. To further determine the relationship between increased Tregs and repressed neutrophils, the frequencies of neutrophils and Tregs in the spleen were analyzed after 150 min of HS exposure, given that the spleen is not only a parenchymatous organ with rich blood flow but also contains many immunocytes. A negative correlation further confirmed the inhibitory effect of Tregs on neutrophils.

Although previous results indicate that increased Tregs may reduce inflammation during HS, heat acclimation involves several physiological changes. Thus, the importance of increased Tregs in protecting individuals from heat stroke is expected. miR-155 is important for the development and function of Tregs. A remarkable reduction in Tregs has been reported in the peripheral immune organs of miR-155-deficient mice^[33]^. As expected, the Treg frequencies in miR-155^-/-^ mice were significantly lower than those in wild-type miR-155^+/+^ mice. Consequently, miR-155^-/-^ mice were tested under the same HS conditions. Surprisingly, the mortality of miR-155^-/-^ mice exposed to HS for 150 min was up to 100%. In contrast, the mortality of Treg-transferred mice exposed to 150 min of HS was as low as 16.7%. Histological scores of the lung and liver also showed a significant protective effect in Treg-transferred mice. This suggests that aggravated inflammatory damage was induced by HS in miR-155^-/-^ mice, and it is also noteworthy that the proinflammatory effects of miR-155 have been shown in other studies^[34, 35]^. This implies a powerful anti-inflammatory effect of Tregs in HS-induced systemic inflammation.

Neutrophils are recognized as the major effector cells in the acute inflammatory response^[36, 37]^. We observed a lower level of neutrophils in the peripheral blood of HA mice during HS-induced systemic inflammation, which prompted us to investigate whether the activation of neutrophils was also suppressed by increased Tregs. A lower concentration of MPO, a major enzyme produced by activated neutrophils, in the supernatant of Treg–neutrophil coculture wells was observed. The immunoregulation of Tregs is related to its epitopes, such as PD-L1, PD-1, and CTLA-4, and the production of inhibitory factors, such as IL-10 and TGF-β^[38, 39]^. In this study, the inhibitory effect of Tregs was reversed after incubation with the PD-L1 antibody but not PD-1, CTLA-4, IL-10, or TGF-β antibodies. The inhibitory effect was abolished when neutrophils were preincubated with the PD-1 antibody. Moreover, pretreatment with 5 μg PD-L1 protein also reduced acute HS-induced mortality in unacclimated mice. Remarkably, there was no protection after pretreating PD-L1^-/-^ mice with Tregs. These results indicate that HS-induced injuries might be regulated by PD-L1 molecules and that neutrophil activation might be suppressed through the PD-1/PD-L1 pathway.

The PD-1/PD-L1 pathway is critical in regulating immune responses^[40, 41]^. PD-L1 plays an essential role in protecting the kidneys and blood–brain barrier from ischemia–reperfusion injury^[42, 43]^. Immunoregulation of Tregs has been shown to involve PD-1/PD-L1, CTLA-4, TGF-β, and IL-10^[44, 45]^. The anti-inflammatory effects of Tregs have been investigated using different approaches. However, our results suggest that the suppressive effect of Tregs on HS-induced neutrophil activation involves only the PD-1/PD-L1 pathway.

We demonstrate for the first time that Treg levels were increased in the spleen and lymph nodes of HA mice. We also uncovered a new protection mechanism of heat acclimation, showing that in HA mice, increased Tregs strongly suppress the HS-induced systemic inflammatory response. Thus, we posit that Tregs may inhibit the activation of neutrophils and improve heat tolerance in mice through the PD-1/PD-L1 pathway. Further in-depth studies on the mechanism of the neutrophil–Treg interaction are warranted. Our results should be significant for devising a potential treatment for systemic inflammatory response syndrome and heatstroke.

## ACKNOWLEDGMENTS

This work was supported by the National Natural Science Foundation of China (grant number 81570497) and the Natural Science Foundation of Chongqing (grant number cstc2020jcyj-bshX0116). We thank Professor Bing Ni for his kind gift GFP-FoxP3 transgenic mice and MS Xiaolang Fu for her technical help.

## ADDITIONAL INFORMATION

Supplementary information accompanies the manuscript on the Experimental & Molecular Medicine website (http://www.nature.com/emm/).

